# Model based analysis of the orderly size-wise activation observed with sinusoidal low frequency alternating current stimulation of peripheral nerves

**DOI:** 10.64898/2026.08.03.742412

**Authors:** Awadh Alhawwash, Ken Yoshida

**Author notes:** Author to whom any correspondence should be addressed.

## Abstract

Extracellular sinusoidal low frequency alternating current (LFAC) stimulation of peripheral motor nerves has been observed to induce size wise activation of nerve fibers, unlike the inverse recruitment order typically seen in extracellular pulsed stimulation. This study aims to explore potential biophysical mechanisms responsible for this phenomenon using computational modeling. Volume conductor model was utilized with a bipolar cuff electrode encasing a single rat-sized fascicle. The extracellular potentials generated by LFAC and pulse stimulation were projected onto the McIntyre–Richardson–Grill models of myelinated motor nerve fibers to examine the activation of fibers ranging from 5.7 to 16μm in diameter. Intracellular and extracellular stimulation were compared for strength-frequency relationships with LFAC (1-20Hz) and strength-duration curves for pulse stimulation. The threshold tracking technique was used to study membrane electrotonus and threshold electrotonus of different fibers to examine subthreshold accommodation in response to LFAC and prolonged pulse stimulation. The simulations revealed that the inverse order of fiber recruitment is an inherent characteristic of extracellular stimulation and is theoretically independent of the stimulation waveform. LFAC showed an inverse strength-frequency relationship (higher frequency, lower threshold current), similar to the inverse strength-duration relationship for pulsed stimulation. Analysis of subthreshold accommodation showed that larger fibers exhibit greater accommodation than smaller fibers, leading to increased activation thresholds as fast Na^+^ activation factor m^3^h decreases while slow K^+^ activation increases, supporting accommodation as a contributor to orderly recruitment. With increasing LFAC frequency (up to 20Hz), these accommodation characteristics were reduced and large-fiber state dynamics shifted toward those of smaller fibers. LFAC was also found to induce subthreshold oscillations that promoted spike initiation during slow depolarization. These findings suggest that LFAC provides a controlled and optimized method for achieving orderly recruitment without the need for complex selective blocking protocols. By leveraging intrinsic membrane properties, LFAC offers a neuromodulation strategy that preserves physiological recruitment order, with direct implications for selective nerve stimulation in clinical and neuroprosthetic applications.

**Author summary:** Electrical stimulation is widely used to activate peripheral nerves in motor rehabilitation and neuroprosthetic devices, but conventional pulse stimulation activates larger nerve fibers first (with lower current intensity), which can induce rapid muscle fatigue and pain. We used well-established and validated computational models of motor nerve fibers (axons) to explore how sinusoidal low frequency alternating current (LFAC) stimulation can produce a more physiological, size-wise recruitment order. We simulated myelinated motor nerve fibers of different diameters individually inside a bipolar cuff electrode and analyzed how the membrane and ion channels changed during stimulation levels that are below thresholds for action potential firing. We found that larger fibers adapt (accommodate) more strongly during the slow depolarization of LFAC: their sodium channels become less open, while potassium activation increases, raising the current required to induce an action potential. Smaller fibers were less affected by this accommodation effect and could reach firing at lower current intensities. We also found that these effects depend on stimulation frequency; at lower frequencies, the accommodation characteristics were more defined (for all fibers), while at higher frequencies they were reduced (for large fibers) and all fiber responses became more similar. Our results suggest that the responses of intrinsic membrane dynamics to LFAC lead fiber recruitment toward a more physiological order, which facilitates the design of safer and more selective nerve stimulation strategies with LFAC.

## Introduction

Electrical stimulation of peripheral nerves (PNS) is widely used in neuromodulation and motor rehabilitation applications. Traditionally, pulsed stimulation has been the dominant method, but recent studies suggest that sinusoidal low frequency alternating current stimulation offers unique features that could improve selectivity and recruitment control to PNS [1, 2]. Unlike conventional extracellular pulsed stimulation, which preferentially activates large-diameter nerve fibers first [3, 4], LFAC has been observed to induce orderly recruitment, where smaller fibers are activated before larger ones [1, 2]. This behavior is particularly desirable for applications such as vagus nerve stimulation (VNS) and functional electrical stimulation (FES), where recruitment order plays a critical role in achieving therapeutic benefit and reducing fatigue [3, 5].

The vagus nerve contains both small-diameter unmyelinated C-fibers and larger-diameter myelinated A- and B-fibers that mediate different physiological effects [6]. VNS applications often require activation of small-diameter fibers, such as in the case of neuromodulation for epilepsy [7], depression [8], and inflammatory disorders [9]. However, conventional pulsed stimulation preferentially activates large-diameter fibers, limiting the ability to selectively target the therapeutic pathways mediated by smaller fibers [3].

Similarly, a major limitation of pulsed stimulation in FES applications is the preferential activation of large, fast-fatiguing motor units, leading to rapid muscle fatigue and discourage patients from using the technology [5]. This recruitment order contrasts with the normal physiological order, the Henneman size principle [10], where smaller, fatigue-resistant Type 1 fibers are recruited first followed by larger fast fatiguing Type 2 motor units.

Our recent studies characterized the frequency dependence and bipolar electrode geometry effects on LFAC stimulation and revealed several key properties [1, 2]. First, the strength-frequency relationship was inversely proportional, where lower stimulation frequencies required higher currents for activation, and vice versa. Second, bipolar cuff electrode geometry influenced activation thresholds, where wider contact separation and greater contact-to-edge distance reducing activation thresholds. Third, both *in-silico* and *in-vivo* results, suggested that LFAC-induced activation follows orderly recruitment only at frequencies below 35 Hz, after which recruitment order becomes inverse. Finally, LFAC was found to produce two activation modes: unitary (single spike per cycle) and burst (multiple spikes per cycle), that were pseudo-random and not synchronized to a fixed point in the sine wave cycle. These activation modes were results from direct cathodic activation under the electrode contacts and virtual activation occurring outside the cuff electrodes [1].

While these findings provided promising insights into LFAC’s electrophysiological properties, the underlying mechanisms responsible for its orderly recruitment feature remain unknown. Traditional extracellular pulse stimulation is widely assumed to invert the recruitment order due to the waveform itself. However, from the nonlinear cable theory, the total current required to excite a fiber is dependent on fiber size and the location of stimulation; whether current is delivered intracellularly (injected into the axoplasm) or extracellularly (applied through a surrounding volume conductor) [11–15]. Theoretically, intracellular stimulation should always induce excitation in the normal order of fiber recruitment; however, with extracellular stimulation several factors play roles in determining excitation thresholds, including fiber size, nerve myelination, electrode location, and electrode geometry [11, 15, 16].

The cable theory can provide an analytical approximation to explain, and to give general insights into, how fiber diameter influence activation thresholds and thus fibers recruitment during intracellular and extracellular stimulation. The general form of the nonlinear cable equation is given by [11–15]:

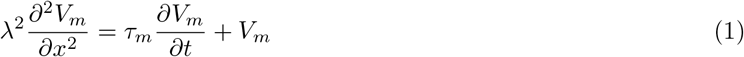

where:

- *τ_m_* is the membrane time constant [s], with *τ_m_* = *R_m_C_m_*.
- *λ* is the space (length) constant [m], with 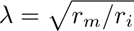
- *R_m_* is the specific membrane resistance [Ω*·*m^2^].
- *C_m_* is the specific membrane capacitance [F/m^2^].
- *r_m_* is the membrane resistance per unit length [Ω*·*m], typically 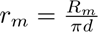 for a cylindrical fiber.
- *r_i_* is the axial (intracellular) resistance per unit length [Ω/m], typically 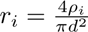 for a cylindrical fiber.
- *ρ_i_* is the intracellular (axoplasmic) resistivity [Ω*·*m].
- *d* is the fiber diameter [m] (often reported in [*µ*m]).
- *R*_in_ is the input resistance at a point of intracellular current injection [11, 12] [Ω], defined as

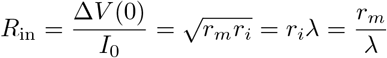

where Δ*V* (0) is the voltage change at the injection site [V] and *I*_0_ is the injected current [A].

During intracellular stimulation, for a cylindrical fiber of diameter *d*, the membrane resistance per unit length, *r_m_*, scales with membrane area as *r_m_ ∝* 1*/d*, while the axial (intracellular) resistance per unit length scales with cross-sectional area as *r_i_ ∝* 1*/d*^2^; therefore, 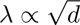 [11, 12]. This proportionality with fiber diameter also scales the input resistance at a point of current injection as 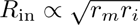 (equivalently *R*_in_ *∝ r_i_λ*) [11, 12], yielding *R*_in_ *∝ d^−^*^3*/*2^ [11]. Thus, to induce a fixed local depolarization of a voltage thresholds *V*_th_, the applied current has to satisfy:

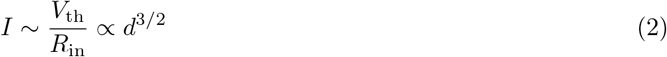

which implies that smaller fibers require less injected intracellular current and therefore favor an orderly (small-to-large) recruitment during intracellular stimulation.

For extracellular stimulation, assuming at steady state (*∂V_m_/∂t* = 0), the cable equation reduces to a purely spatial relationship between the induced transmembrane polarization and the extracellular potential (*V_e_*) distribution, where that extracellular excitation becomes governed by the spatial variation of *V_e_*(*x*) along the fiber (i.e., the activating-function term [13, 14]) and influenced by the cable properties through the length constant *λ* [11]. However, for a nerve trunk/bundle containing different nerve fibers, the applied extracellular current does not flow equally within bundle’s fibers where the axial resistance, *r_i_*, dominantly influences each fiber’s activation thresholds, which scales as 1*/d*^2^. Thus, the excitation current threshold with extracellular stimulation can be approximated to scale as factor for both the circumference changes (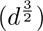) and axial resistance (*d^−^*^2^) that are directly related to fiber diameter [11]. Under simplifying assumptions (e.g., comparable *V_e_*(*x*) along fibers and current partition dominated by the axial pathway), the extracellular threshold current can be approximated to scale as *I*_th,extra_ *∝ d*^3*/*2^ *·* 1*/d*^2^ *∝ d^−^*^2^ predicting preferential extracellular activation of larger fibers (inverse recruitment) [11]. Furthermore, this arises the possibility that extracellular stimulation inherently favors large fiber activation, and that pulses amplify this effect due to the induced rapid depolarization rate, 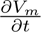 from the cable equation. Higher sine wave frequencies (35Hz to 1kHz based on our previous study) were found to induce inverse recruitment order, likely due to the first derivative of the current waveform increasing with frequency (f), such that 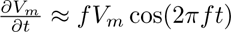, which preferentially excited large fibers [1]. These factors of membrane morphology and time-dependent changes provide analytical insights as of why pulse stimulation and sinusoidal high frequency stimulation induce inverse order of fiber recruitment. In terms of LFAC extracellular stimulation, these factors could contribute, in part, to the orderly recruitment feature given that the depolarization rates, 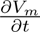 of membrane potential would be expected to be more gradual and slow comparing to pulse and high frequency stimulation.

Another possible mechanism is that LFAC could induce subthreshold nerve accommodation, thereby shaping the recruitment order in a fiber-dependent manner. Subthreshold accommodation, described as a reduction in excitability (increased threshold) following subthreshold membrane depolarization, was found to impact subthreshold prolonged pulse stimulation and influence different excitability patterns when characterizing axonal excitation [17, 18]. Bostock and others studied accommodation mechanisms, which were attributed primarily to the dynamics of voltage-gated ion channels; where sodium channels transition to their inactivated state while potassium channels activate, reducing the probability of action potential firing during slow changes in membrane potential [17, 18]. Studies comparing the accommodation effect of different nerve fiber sizes suggest that large fibers exhibit accommodation more than small fibers when tested with ramp or exponential stimuli [19–23]. Given the nature of LFAC waveform, it is possible that accommodation would contribute to induce orderly fiber recruitment where large fibers would accommodate to slowly changing current more than small fibers.

To investigate these mechanisms, this study presents a model-based analysis using finite element models (FEM) and the McIntyre-Richardson-Grill (MRG) models of myelinated motor nerve fibers [24]. Specifically, we aim to: 1) compare LFAC and pulsed stimulation with bipolar cuff electrode to determine whether recruitment order inversion is an inherent feature of extracellular stimulation or a consequence of waveform properties. 2) Examine intracellular stimulation via current clamp experiments, to show that both LFAC and pulses produce orderly recruitment when applied intracellularly. 3) Conduct accommodation experiments using Bostock’s threshold tracking technique [17, 18], to investigate fiber-specific accommodation to LFAC vs. long-duration pulses. 4) Analyze ion channel dynamics to explore their contribution to fiber-dependent activation.

## Methods

Computer simulations were performed to examine the dynamics of sinusoidal LFAC stimulation with specific cuff electrode geometries as shown in Fig 1B. The experiments mostly followed the approach of *in-silico* part in (Alhawwash et. al. 2025 [1]). Briefly, the models consisted of 1) inhomogeneous volume conductor models of the rat tibial nerve, used to approximate the extracellular potentials (*V_e_*). 2) The MRG myelinated motor nerve fiber models [24] to approximate the nerve’s response to extracellular (*V_e_*) or intracellular stimulation. To examine intracellular stimulation, the MRG myelinated motor nerve fiber models were used with an injected current without applying an extracellular stimulation.

**Fig 1.**
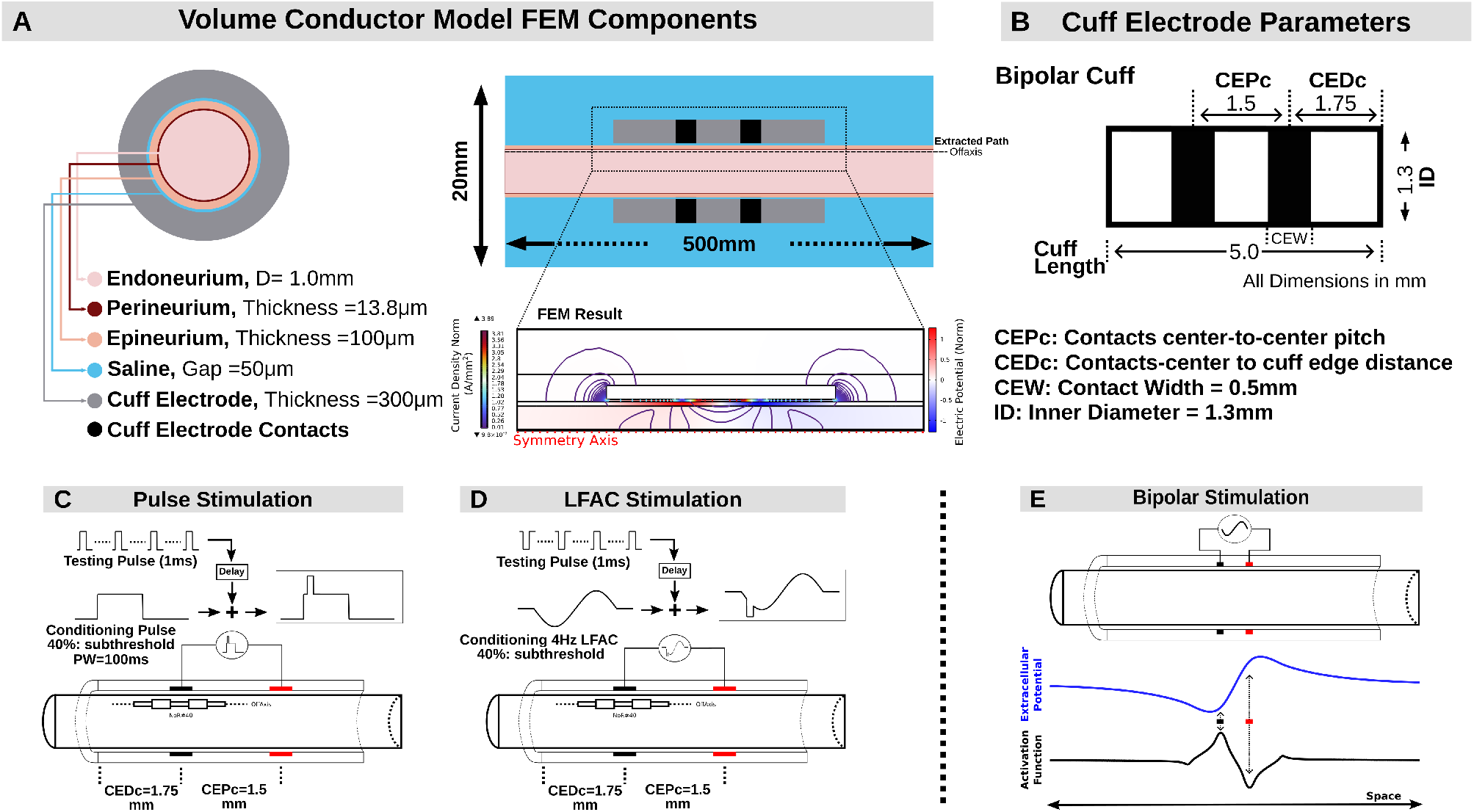
Volume conductor model and cuff electrode configurations. (A) Cross-sectional and longitudinal views of the volume conductor model, showing the finite element model (FEM) components and the bipolar cuff electrode. The FEM simulation of the cuff electrode configuration illustrates the extracellular potential distribution with 4Hz LFAC stimulation at the peak time of the sinusoidal waveform. The potential distribution was extracted along a linear path, 1*µ*m from the inner perineurium wall, illustrated as Offaxis. (B) Schematic representation of the bipolar cuff electrode parameters used in this study; including contact width (CEW) that was set to 0.5mm, inner diameter (ID) of 1.3 mm, contact separation distance -contacts center-to-center pitch (CEPc), contact-center to cuff edge distance (CEDc), and cuff electrode length. (C-D) Experimental setup for subthreshold accommodation testing using threshold tracking. The bipolar cuff electrode was used to apply a conditioning stimulus, followed by a delayed 1ms testing pulse to track threshold changes over time. (C) Pulse stimulation: a 100ms subthreshold conditioning pulse (40%of activation threshold) was applied, followed by a series of delayed 1ms testing pulses. (D) LFAC stimulation: a single cycle of 4 Hz LFAC set at 40% of activation threshold was used as the conditioning stimulus. The testing pulse polarity followed the LFAC polarity at each time point. The same testing protocol was applied for both conditions. (E) Extracellular potential (blue) and activation function (black) of typical profile for bipolar stimulation configuration with inset marks of contacts locations. NoR: Node of Ranvier. All drawings are not to scale.

### Volume Conductor Model (FEM Simulations)

2D axial-symmetric finite element models of the rat tibial nerve bundle were created in COMSOL Multiphysics (V6.2, Comsol Inc, Burlington, MA). The modeling methods followed the steps and conditions reported in the *in-silico* part of (Alhawwash et. al. 2025 [1]) with major differences as follows: Bipolar cuff electrode, with constants inner diameter of 1.3*µ*m and thickness of 300*µ*m, were modeled with dimensions shown in Fig 1B and positioned centrally around the nerve with 50*µ*m gap.

The electrical properties of the model’s components are listed in Table 1 and their dimensions are listed in Fig 1A. Boundary conditions, infinite element domains, and meshing were performed as in [1]. Sinusoidal and monophasic rectangle pulses were applied as normal current densities and were delivered through each contact of the bipolar electrode. The cathodic and anodic magnitudes were set to 1 A*_p_* per unit contact area. In all settings, the contacts were acting as current sources. LFAC stimulation was tested with frequencies of 1, 2, 3, 4, 8, and 20 Hz. Pulse stimulation waveform was generated with a rising time of 20*µ*s, a frequency of 1Hz, and pulse widths of 0.05, 0.1, 0.2, 0.5, 1, 2, 5, 10, and 100 ms. Time-dependent studies were used to solve for electric potentials with simulation runtime equal to 2 periods of each sinusoidal frequency or 300ms for pulse stimulation. Potential distribution (*V_e_*) was extracted along the longitudinal Offaxis path located 1*µ*m from the inner perineurium wall as shown in Fig 1A. The extracted *V_e_* was then linearly interpolated in space to ensure a consistent spatial resolution of 5*µ*m between sample points.

**Table 1.** Electrical properties of the volume conductor model.

| Material | Conductivity<br>(S/m) | Relative<br>Permittivity | References |
| --- | --- | --- | --- |
| Saline | 1.76 | 80 | [25, 26] |
| Endoneurium (radial) | 0.083 | 80 | [27] |
| Endoneurium (longitudinal) | 0.57 | 80 | [27] |
| Perineurium | $2.7 \times 10^{-4}$ | 2018 | [28] |
| Epineurium | 0.018 | $9.4 \times 10^6$ | [28] |

### Myelinated Active Nerve Models (NEURON Simulations)

#### Extracellular Stimulation

The McIntyre-Richardson-Grill (MRG) myelinated nerve fiber models [24] were utilized in this study following the approach described in [1]. However, the simulation paradigms consisted of extracellular stimulation (using *V_e_* from the FEM) and intracellular stimulation. Simulations were performed using NEURON V8.0 [29], utilizing Python (V3.9, Python Software Foundation, Wilmington, DE) and MATLAB (R2023b, MathWorks, Natick, MA). In both settings, seven fiber diameters were simulated (5.7, 7.3, 8.7, 10, 11.5, 14, and 16*µ*m) with 80 nodes of Ranvier (NoR) to investigate the recruitment properties of LFAC and pulse stimulation within and around the range of rat tibial nerve axons [30, 31]. For each fiber, we collected the strength-frequency curves during LFAC stimulation and strength-duration curves during pulse stimulation.

The extracellular potential distribution (*V_e_*) extracted from the FEM model was spatially aligned with the middle node of Ranvier (Node 40). The location of the first electrode contact, initially set at *x* = 0, was shifted to align with Node 40. The position of this contact was identified using the activation function, defined as the second spatial derivative of *V_e_*, evaluated at the sinusoidal peak time or at the midpoint of the pulse waveform. The potential distribution *V_e_*was spatially interpolated at the centers of each axonal MRG compartment, including paranodal myelin attachment segments (MYSA), paranodal main segments (FLUT), and internodal segments (STIN). The interpolated *V_e_* was then scaled by a peak testing amplitude with cathodic phase being applied at the first contact initially.

Using the extracellular mechanism in NEURON, *V_e_* was applied to the axon models, and simulations were executed for 1.5 cycles of each sinusoidal frequency with a time step of 0.01ms, employing NEURON’s backward Euler integration method. For pulse stimulation, simulations were conducted for 300ms. Evoked action potentials (APs) were tracked and a binary search method was used to determine activation thresholds. Since LFAC produced both cathodic and virtual activations, the site of AP initiation was recorded relative to the electrode contacts and stimulation phase. Activations occurring within the cuff were classified as direct cathodic activations, while those initiated outside the cuff were identified as virtual cathodic activations, henceforth called “virtual activations.”

Subthreshold accommodation was investigated using Bostock’s threshold tracking technique [17, 18]. The bipolar cuff electrode delivered a conditioning stimulus followed by a delayed 1ms test pulse to track threshold changes over time. For pulse stimulation, a 100ms subthreshold conditioning pulse, set at 40% of the activation threshold, was applied, followed by a series of delayed 1ms test pulses (Fig 1C). For LFAC stimulation, a single cycle of 4Hz LFAC at 40% of the activation threshold was used as the conditioning stimulus (Fig 1D), with the test pulse polarity following the LFAC polarity at each time point.

Initially, each conditioning stimulus was applied to determine the activation threshold for each of the seven fiber diameters. During threshold tracking experiments, each fiber was tested using its corresponding 40% activation threshold for the conditioning stimulus. Electrotonus, defined as the membrane potential at 40% threshold, and activation threshold changes as a function of fiber diameter were analyzed. Subsequently, threshold reduction over time was calculated as the percentage change in test pulse threshold relative to baseline (without conditioning stimulus). In both setting, the conditioning stimulus was applied at a simulation time of 50ms to ensure steady-state conditions before threshold tracking begins.

For LFAC activation dynamics analysis, the response to a 4Hz sinusoidal LFAC stimulus was investigated by tracking the nodal membrane potential, transmembrane ionic currents, channel gating variables, and gating time constants at all nodes of Ranvier. We looked at the mechanisms of the nodal membrane dynamics which are governed by fast sodium Na^+^, persistent sodium Na^+^, slow potassium K^+^ channels, and leakage mechanisms in parallel with the nodal capacitance. Furthermore, we computed the dimensionless gating factors for the sodium channels, including the fast Na^+^ activation factor *m*^3^*h* and the persistent Na^+^ activation factor *p*^3^, to assess diameter-dependent changes via overall probability of channels activation. Three fiber diameters were evaluated (7.3, 10, and 16*µ*m) under two subthreshold stimulus protocols: (1) a *fiber-specific* condition, in which each fiber was stimulated at its own subthreshold peak current; and (2) a *common-amplitude* condition, in which all fibers received the same peak current set to the subthreshold level of the 7.3*µ*m (230.9*µ*Ap). In both protocols, the “subthreshold” level was defined as the largest LFAC amplitude that elicited sustained subthreshold responses without action potential initiation over the analysis window of three LFAC cycles, in order to compare diameter-dependent channel state evolution under LFAC stimulation.

To evaluate the frequency dependence of these dynamics, the analysis was repeated at 1, 2, 3, 8, and 20Hz using only the *fiber-specific* protocol. For each frequency, the same set of outputs was recorded as for 4Hz (membrane potential and gating variables, sodium-channel gating factors, membrane current components, and gating time constants) over three LFAC cycles.

#### Intracellular Stimulation

Intracellular stimulation was explored to provide computational examples of nerve responses to injected current into a single node of a cable model. Simulations were performed in NEURON utilizing the IClamp (current clamp) mechanism, which injects current as an additional current source within the model. In this case, the classical definition of a current clamp does not strictly apply, as the injected current does not represent a transmembrane current but is directly applied inside the cell. For pulse stimulation, strength-duration curves were acquired for the seven fiber diameters using the NEURON IClamp mechanism [29], where pulse width and amplitude were systematically varied. For LFAC stimulation, sinusoidal waveforms were generated and applied to the IClamp mechanism using the Izap function [32]. Each LFAC simulation was conducted for two cycles per applied frequency. As with extracellular stimulation, evoked action potentials were tracked using a binary search method to determine activation thresholds.

## Results

### Intracellular versus extracellular stimulation

Intracellular stimulation was examined using a current clamp approach in NEURON, applying either pulsed stimulation or LFAC at different frequencies to the seven fibers. Activation thresholds were acquired for each fiber for both waveforms, revealing an orderly recruitment pattern; where smaller fibers required lower activation currents than larger fibers as seen in Fig 2A and Fig 2B. For pulsed intracellular stimulation, Fig 2A, the activation threshold followed the classical strength-duration relationship, where longer pulse durations led to a reduction in the required activation current. Similarly, for LFAC intracellular stimulation, Fig 2B, the strength-frequency relationship was revealed for the explored frequency range (1-20Hz); where higher LFAC frequencies required lower current amplitudes to reach threshold. The results indicate that both pulsed and LFAC stimulation applied intracellularly maintain the physiological recruitment order (size-wise) of nerve fibers. However, pulse stimulation required higher activation thresholds within the range of 0.2-3.6 *µ*Ap while LFAC required lower activation thresholds below 1.2 *µ*Ap.

**Fig 2.**
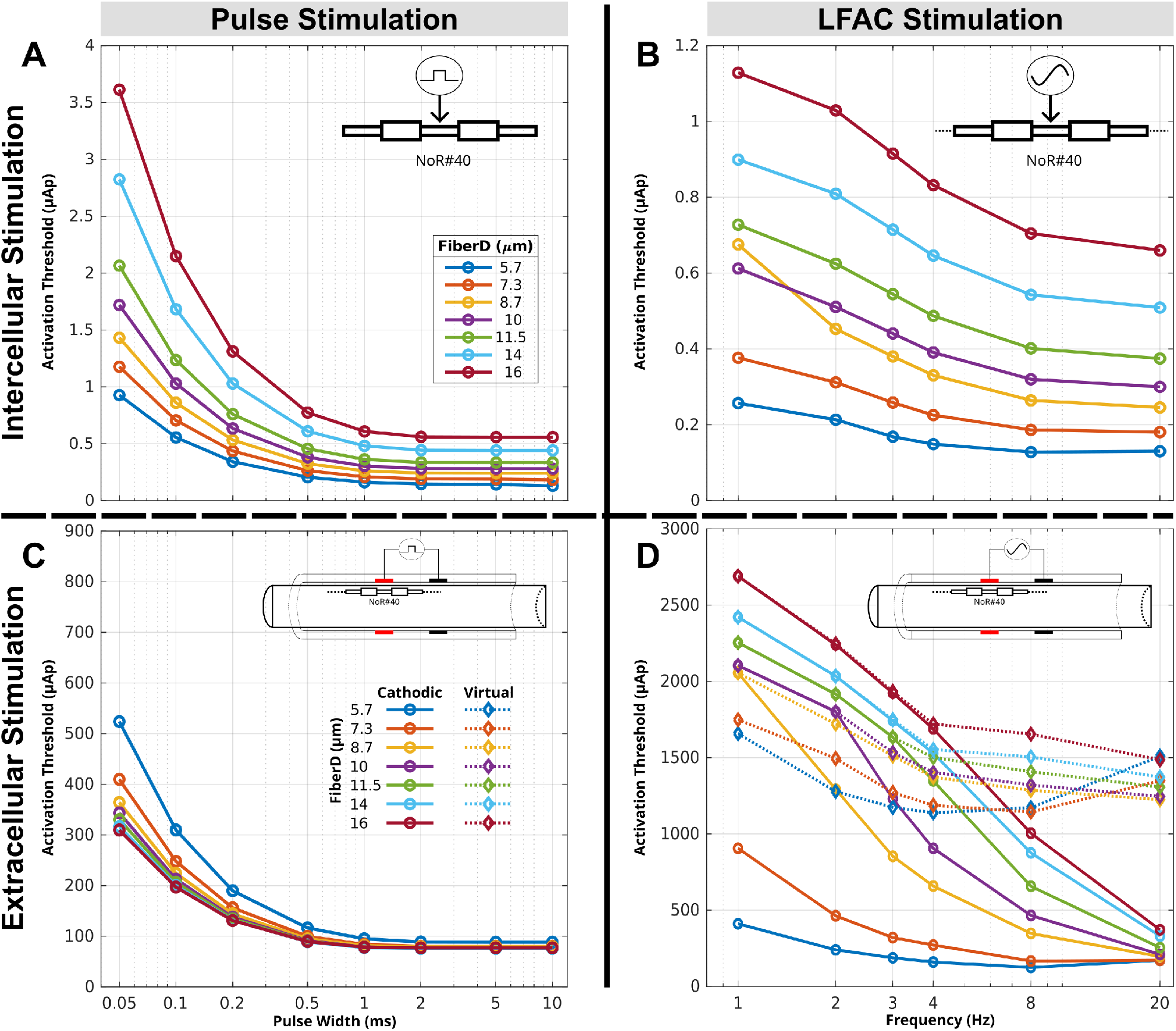
Comparison of intracellular and extracellular stimulation thresholds using LFAC and pulse waveforms. (A, B) Intracellular stimulation: Activation thresholds for pulse stimulation (A) and LFAC stimulation (B) applied via current clamp in NEURON across seven fiber diameters (5.7-16 *µ*m). Both LFAC and pulse stimulation revealed orderly recruitment, where smaller fibers activate at lower thresholds than larger fibers. Pulse stimulation follows the classical strength-duration relationship, while LFAC thresholds decrease as frequency increases. (C, D) Extracellular stimulation: Activation thresholds for pulse stimulation (C) and LFAC stimulation (D) applied through a bipolar cuff electrode of the same geometry. Pulse stimulation inverts the recruitment order, preferentially activating larger fibers with only cathodic activation mode. In contrast, LFAC preserves orderly recruitment with cathodic activation mode and shows frequency-dependent recruitment order with the virtual mode. NoR: Node of Ranvier. Cathodic legends in C are for both C and D. Virtual legends are only for D (LFAC case).

In contrast to intracellular stimulation, extracellular stimulation showed different recruitment behaviors between pulse and LFAC waveforms. Pulsed stimulation showed the classical behavior of preferentially activating larger fibers at lower thresholds, within the range of 309-523 *µ*Ap for 0.05ms pulse width to 75-88 *µ*Ap for 10ms pulse width. Regardless of the pulse width, the recruitment order was inverted, where larger fibers were activated before smaller fibers, as shown in Fig 2C. For LFAC extracellular stimulation, Fig 2D, LFAC produced cathodic and virtual activations. Direct cathodic activation occurred directly under the contact when it acted as a cathode. The strength-frequency relationship was maintained, where strength and frequency are seemingly inversely proportional, with higher frequencies requiring lower activation currents. However, compared to intracellular stimulation, direct cathodic activation required substantially higher current amplitudes within the range from 411-2689 *µ*Ap at 1 Hz to 171-371 *µ*Ap at 20 Hz. The orderly recruitment pattern was preserved, with smaller fibers being activated at lower thresholds than larger fibers.

On the other hand, virtual activations occurred outside the cuff electrode and their thresholds in Fig 2D show more fiber-frequency dependent behavior. Overall, virtual activation thresholds were higher than cathodic with maintained strength-frequency relationship except at 20 Hz; where 5.7 and 7.3 *µ*m fibers required higher currents. Besides those two fiber sizes, the the recruitment order was normal, small to large.

### Subthreshold Accommodation to 4Hz LFAC and 100ms pulse

Examining subthreshold accommodation revealed that both pulse and LFAC stimulation induce fiber dependent reduction in activation thresholds. Fig 3 shows side-by-side comparison of subthreshold accommodation induced by 4Hz LFAC and a 100ms conditioning pulse. The effects of subthreshold stimulation on threshold were examined by comparing electrotonus (membrane potential at 40% threshold of each fiber), threshold reduction over time, and activation threshold changes as a function of fiber diameter. Fig 3A and Fig 3B shows the electrotonus response at Node 40 for the seven fibers with a subthreshold conditioning stimuli. Both pulse and LFAC show more depolarization as fiber diameter increased with differences from resting membrane potential. Since the conditioning stimuli thresholds were higher for LFAC comparing to the pulse, the depolarized membrane potential magnitude with LFAC reached higher values than pulse conditioning. Pulse stimulation produced rapid overshoot depolarization (with fibers greater than 8.7*µ*m) followed by gradual decrease, whereas smaller fibers induced slower depolarization followed by gradual decrease. On the other hand, LFAC resulted in oscillatory depolarization and hyperpolarization phases.

**Fig 3.**
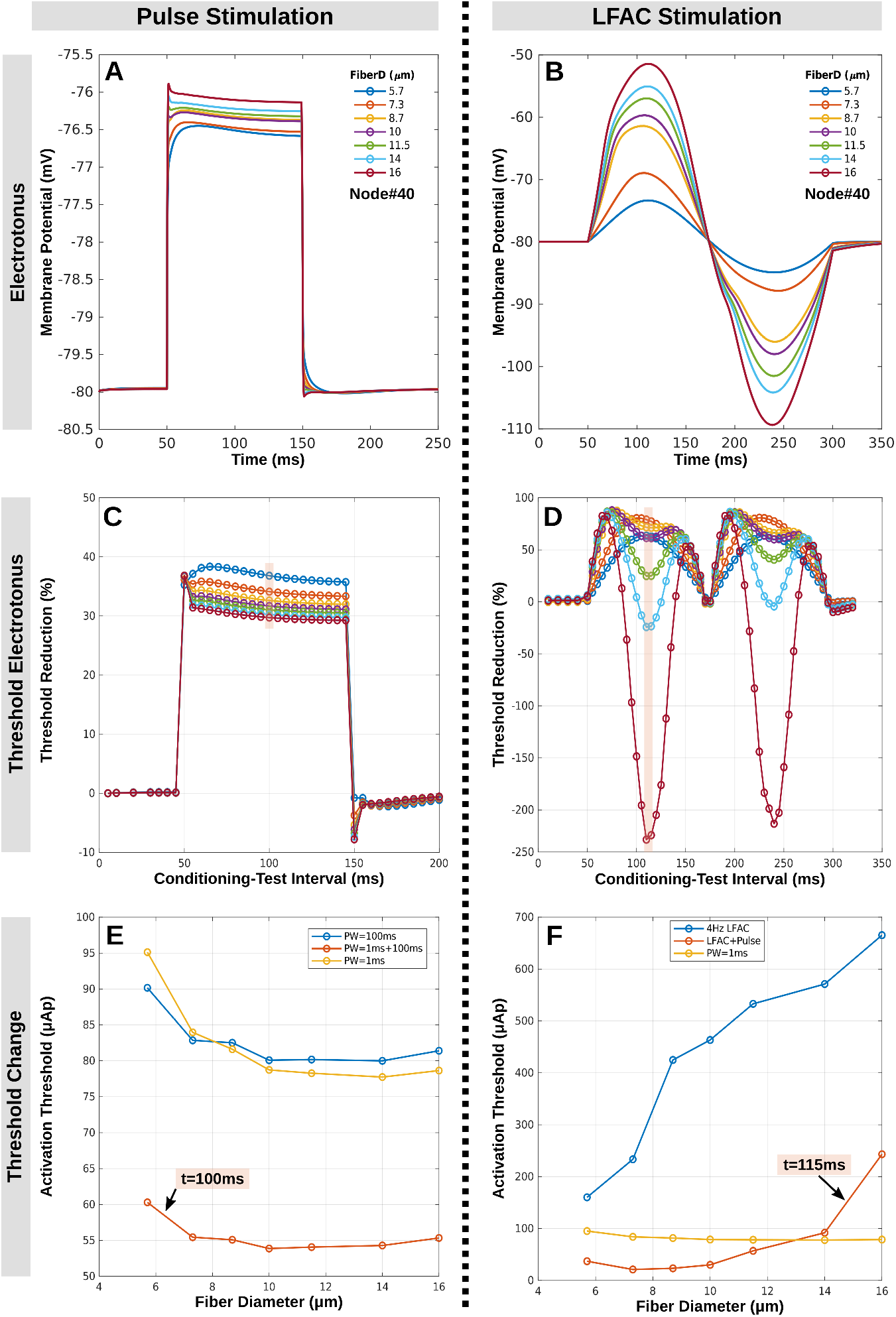
Comparison of subthreshold accommodation induced by 4Hz LFAC and a 100ms conditioning pulse. (A, B) Membrane potential response at Node 40 for different fiber diameters with a 100ms subthreshold conditioning pulse (A) and 4Hz LFAC (B). (C, D) Threshold reduction over time, measured as the percentage change in test pulse threshold compared to baseline, for pulse (C) and LFAC (D) conditions. (E, F) Activation threshold as a function of fiber diameter at selected time points. (E) The conditioning pulse reduced activation thresholds for larger fibers more than smaller fibers, whereas (F) LFAC preserved orderly recruitment, with small fibers maintaining lower thresholds than large fibers. Highlighted regions in C and D illustrate examples of activation thresholds at specific time points (100ms for pulse and 115ms for LFAC in E and F).

In Fig 3C and Fig 3D, the testing pulse threshold reduction over time, measured as the percentage change in test pulse threshold compared to baseline are shown for pulse and LFAC conditions. In Fig 3C, the conditioning pulse resulted in reduced thresholds for all fibers following the depolarization behavior of the electrotonus over time with small fibers showing higher reduction than large fibers. The pulse activation thresholds as a function of fiber diameter at representative time points are shown in Fig 3E along with the conditioning stimulus’ thresholds, which shows inverse order of recruitment with and without the conditioning pulse.

On the other hand, in Fig 3D, the LFAC conditioning resulted in time-fiber-dependent threshold changes that vary cyclically, with significant modulation observed for larger fibers around the peak times in both phases of the LFAC waveform. The pulse activation thresholds as a function of fiber diameter at representative time points are shown in Fig 3F along with the conditioning LFAC thresholds, which shows orderly recruitment with the LFAC conditioning and inverse without it. Panels of Fig 4 expanded those in Fig 3F to illustrate how fibers recruitment order was modulation for each time delay of the test pulse during the single LFAC cycle. A supplementary video (S1 Video) illustrates the temporal evolution of these threshold curves across the LFAC cycle.

**Fig 4.**
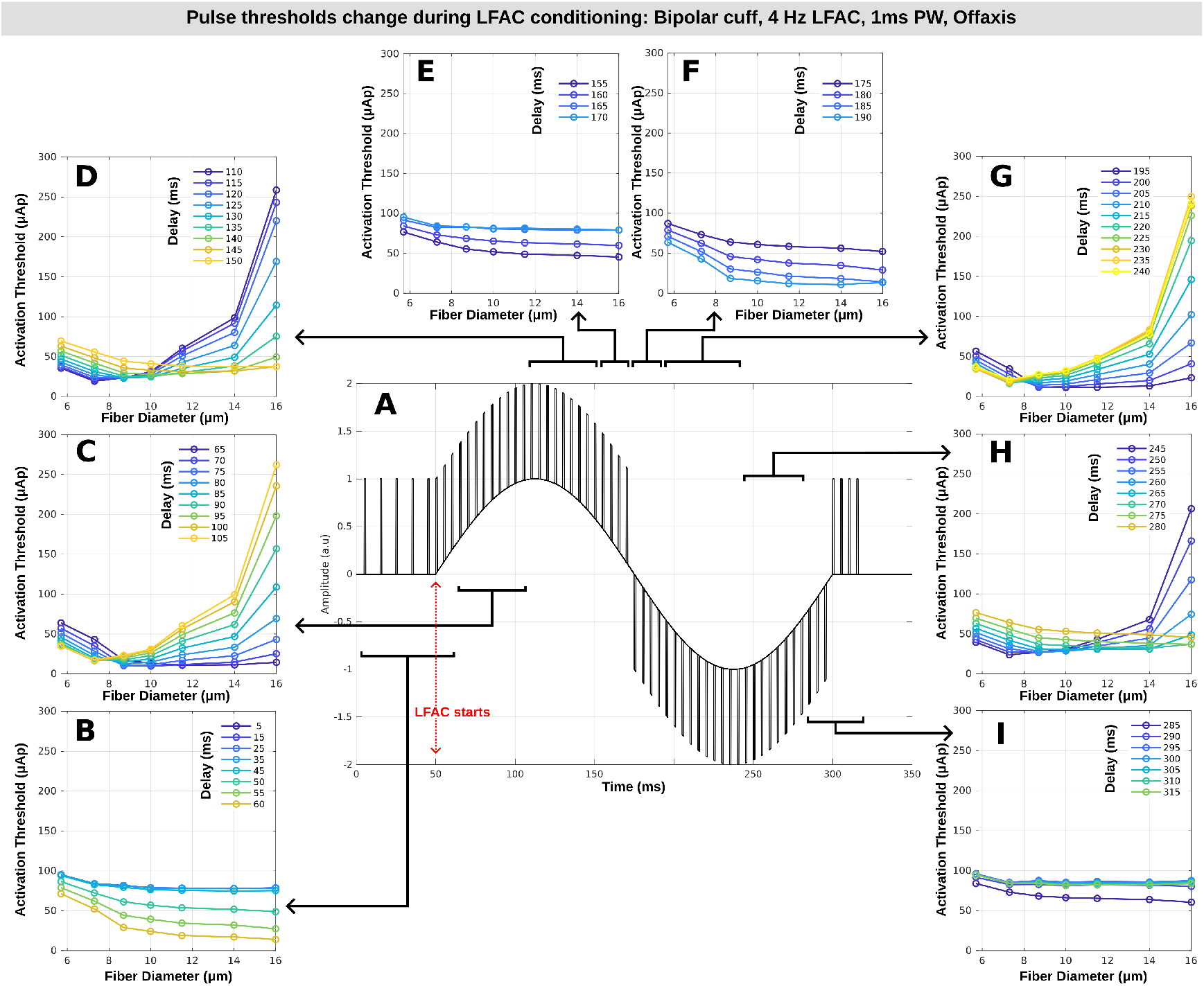
Pulse activation thresholds dynamically modulated by sinusoidal LFAC stimulation across fiber diameters and time. (A) A 4Hz sinusoidal LFAC waveform is shown, with vertical bars marking pulse timing delays relative to LFAC onset. Panels (B to I) display activation thresholds for 1ms pulses delivered at sequential time delays across the LFAC cycle. Each curve represents thresholds across fiber diameters (5.7 to 16*µ*m) at a given delay. For each fiber, LFAC amplitude was set to 40% of its corresponding LFAC activation threshold. A supplementary video (S1 Video) illustrates the temporal evolution of these threshold curves across the LFAC cycle.

### LFAC activation dynamics

By focusing on the direct cathodic activation of LFAC, Fig 5 shows representative examples of 7.3*µ*m and 16*µ*m fibers membrane potential responses to a single cycle of 4Hz LFAC stimulation (with 50ms delay) at different current levels. In Fig 5A and Fig 5B, at 90% of the activation threshold of 7.3*µ*m fiber (230.9*µ*Ap), both 7.3*µ*m (A) and 16*µ*m (B) fibers were not activated, but 7.3*µ*m fiber shows subthreshold oscillations, but no action potential was generated. In Fig 5C and Fig 5D, at the activation threshold of the 7.3*µ*m fiber (258 *µ*Ap), the 7.3*µ*m fiber is activated while the 16*µ*m fiber remains below threshold. As shown in the temporal traces, the AP initiating was at the top of the observed subthreshold oscillations. With further current increase, Fig 5E and Fig 5F, at the activation threshold of the 16*µ*m fiber (1689 *µ*Ap), both fibers are activated, but two activation behaviors are observed for 7.3*µ*m fiber; the direct cathodic activation under the cathodic electrode contact and the virtual activation outside the cuff electrode. At this current level, it was well above the virtual activation thresholds for 7.3*µ*m fiber (Fig 2D); however it was the minimum current needed to induce direct cathodic activation for 16*µ*m fiber.

**Fig 5.**
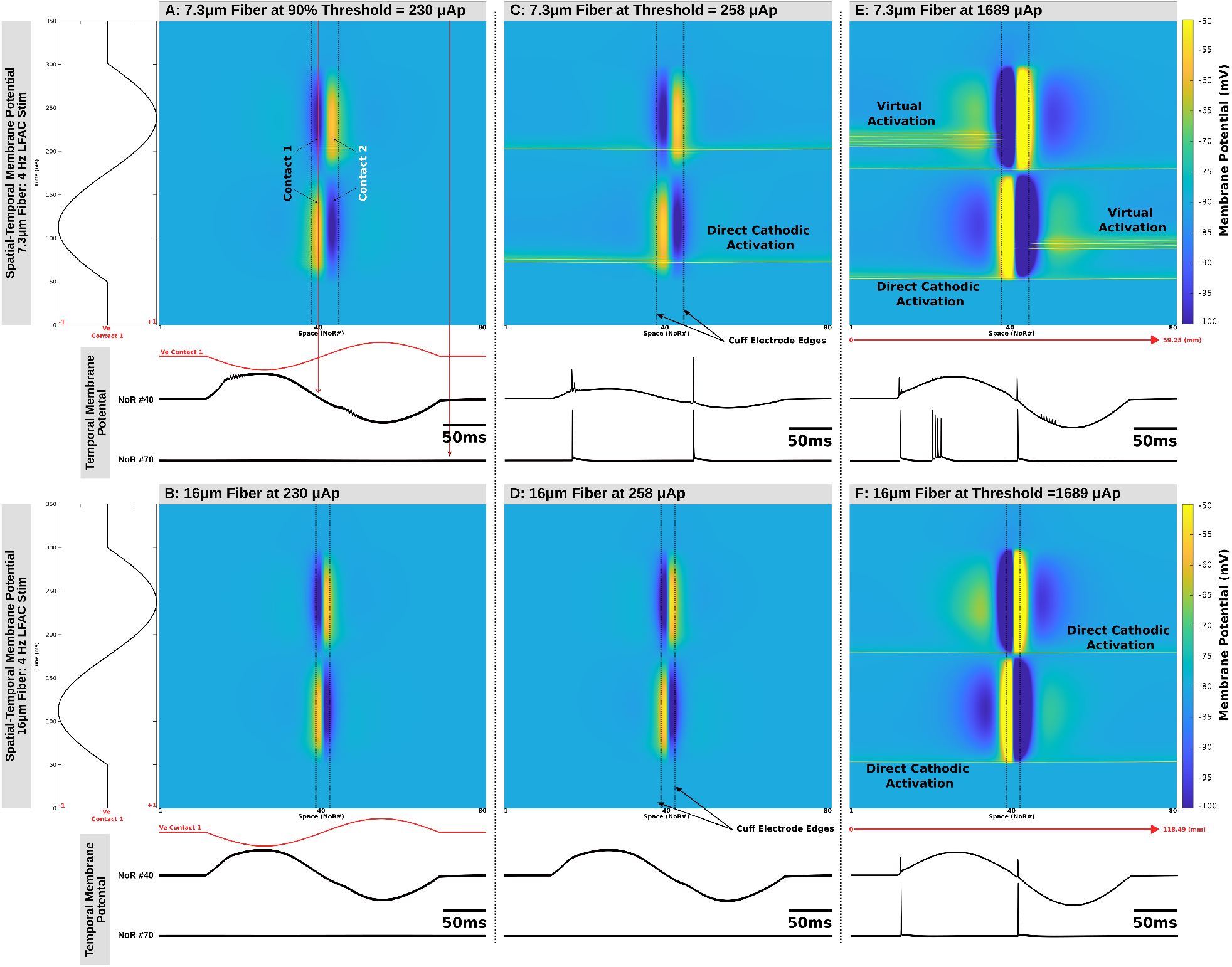
Spatial-temporal behavior of membrane potential in response to LFAC stimulation at different current levels for 7.3*µ*m and 16*µ*m fibers. LFAC was applied through a bipolar cuff electrode, and the membrane potential was examined over a single cycle with a 50 ms delay. (A, B) At 90% of the activation threshold of the 7.3*µ*m fiber, both 7.3*µ*m (A) and 16*µ*m (B) fibers were not activated, but 7.3*µ*m fiber shows subthreshold oscillations. (C, D) At the activation threshold of the 7.3*µ*m fiber, the 7.3*µ*m fiber was activated while the 16*µ*m fiber remained below threshold. (E, F) At the activation threshold of the 16*µ*m fiber, both fibers were activated with two activation modes for 7.3*µ*m: direct cathodic activation under the cathodic electrode contact and virtual activation outside the cuff electrode. Dotted vertical lines mark cuff electrode edges and node positions are normalized to node numbers, but the actual spatial resolution depends on node-to-node distances (750*µ*m for the 7.3*µ*m fiber and 1500*µ*m for the 16*µ*m fiber, total lengths are shown in E and F). Below each surface plot, temporal membrane potential traces from Node 40 and Node 70 illustrate activation timing and oscillatory behavior. NoRs: Nodes of Ranvier. NoRs locations are not to scale.

To investigate the dynamics of cathodic LFAC activation and the mechanisms underlying orderly recruitment, we applied three cycles of 4Hz LFAC with a 50 ms delay to three different fibers (7.3*µ*m, 10*µ*m, and 16*µ*m) and analyzed their gating variables, ionic currents, and time constants at subthreshold levels using two stimulus protocols. Since ion channels are only located in the nodes of Ranvier, we looked at the temporal changes of the channels’ variables at node 40; the center of the models where it was the location of initiating direct cathodic activation underneath one of the electrode contacts.

During the *fiber-specific* condition, each fiber’s input current was set to its own subthreshold LFAC current (7.3 *µ*m: 230.9*µ*Ap; 10 *µ*m: 434.6*µ*Ap; 16 *µ*m: 641.8*µ*Ap). Fig 6 summarizes membrane potential responses together with gating variables and gating factors during three LFAC cycles, while Fig 7 and Fig 8 report the corresponding ionic/membrane currents and channels gating time constants. Fig 6A shows the membrane potential responses for the three fibers, which reveals that there are subthreshold oscillations during the depolarizing phase of LFAC. In Fig 6A, three regions (I, II, and III) are highlighted and zoomed in to clearly show the magnitude and the oscillatory behavior of the membrane potential. With respect to the sinusoidal phase, membrane potential and gating variables followed depolarizing behavior during the cathodic current phase and hyperpolarizing behavior during the anodic current phase. The subthreshold oscillations occurred during the the rising (depolarizing) phase of LFAC; however, when the second contact acts as a cathode, those oscillations are propagated to be seen during the hyperpolarizing phase. During the time of region I and III, node 40 was under depolarizing portion of the waveform and hyperpolarizing portion during time of region II. Despite comparable near-threshold operating conditions, the magnitude values of membrane potential responses exhibit diameter-dependent subthreshold behavior, providing the basis for mechanistic comparison of channel state evolution.

**Fig 6.**
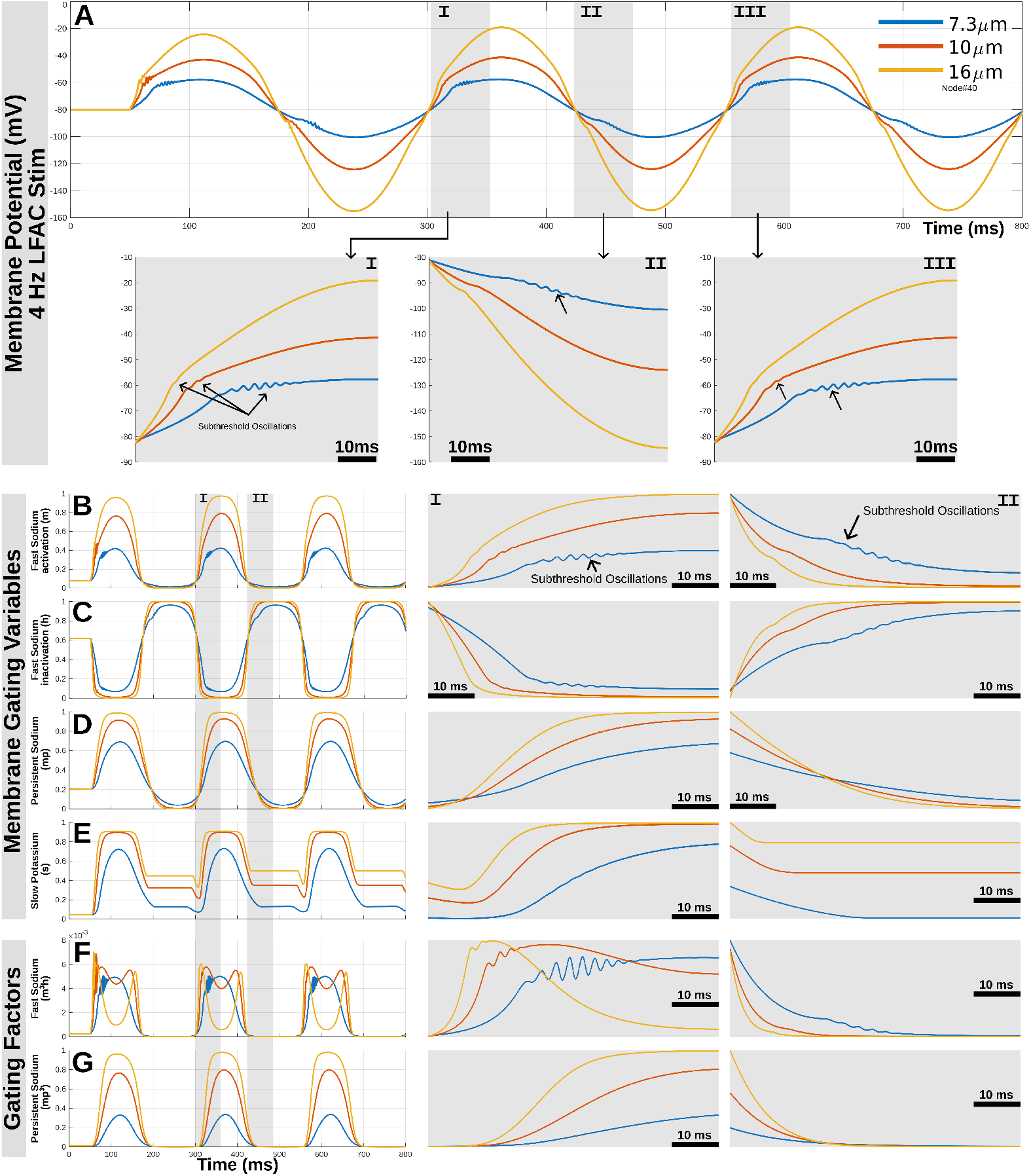
Subthreshold membrane and channel-state dynamics across fiber diameters under LFAC stimulation during the *fiber-specific* condition. Three cycles of 4Hz LFAC were applied following a 50ms resting period, where each fiber was driven at its own subthreshold peak LFAC current (7.3 *µ*m: 230.9*µ*Ap; 10 *µ*m: 434.6*µ*Ap; 16 *µ*m: 641.8*µ*Ap). (A) Membrane potential at Node 40 (model center; location of the cathode-first contact) for 7.3*µ*m (blue), 10*µ*m (orange), and 16*µ*m (yellow) fibers, showing subthreshold oscillations near the depolarizing portion of the LFAC cycle. (B–E) Diameter-dependent gating-variable responses for fast Na^+^ activation (*m*), fast Na^+^ inactivation (*h*), persistent Na^+^ activation (*mp*), and slow K^+^ activation (*s*), with oscillatory features most prominent in the fast Na^+^ variables. (F) Fast Na^+^ activation factor *m*^3^*h*, showing a reduction in the larger fibers as depolarization progresses. (G) Persistent Na^+^ activation factor *mp*^3^, which reflects no oscillations and increases in magnitude with fiber diameter only during depolarization. Shaded regions (I, II, and III) indicate segments of the second and third LFAC cycles that are shown as zoomed views to highlight fiber-specific differences during depolarization (I and III) and hyperpolarization (II).

**Fig 7.**
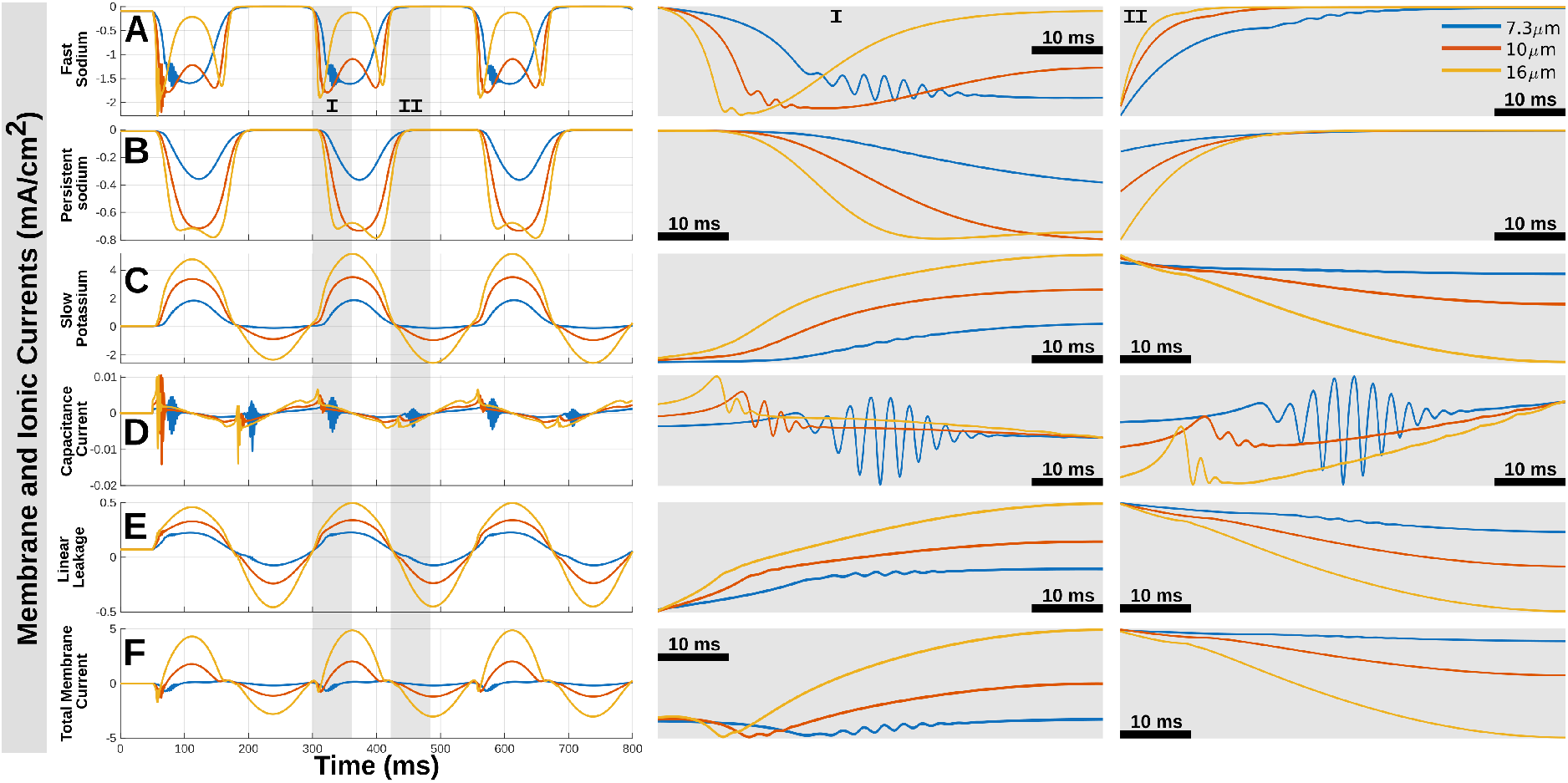
Membrane current densities across fiber diameters under subthreshold LFAC stimulation during the *fiber-specific* condition. Using the same stimulation protocol as in Fig 6, this figure shows the subthreshold behavior of individual ionic, capacitive, and leak current components at Node 40 for the three fibers: 7.3*µ*m (blue), 10*µ*m (orange), and 16*µ*m (yellow). In all panels, inward current densities are shown as negative values and outward current densities are shown as positive values. (A–C) Ionic currents of fast Na^+^ (A), persistent Na^+^ (B), and slow K^+^ (C) showing diameter-dependent modulation over the LFAC cycles, with the most changes occurring during the depolarizing portion of the waveform. The inward fast Na^+^ current increases transiently and then attenuates more rapidly in the larger fibers, together with subthreshold oscillations around depolarization. The persistent Na^+^ current shows a similar but slower modulation, most evident in the 16*µ*m fiber. (D) Capacitive current, which reflects the high-frequency subthreshold oscillations most clearly, consistent with rapid voltage fluctuations in the near-threshold regime. (E) Linear leak current, whose magnitude follows the LFAC-driven membrane potential and varies across diameters. (F) Total membrane current, showing oscillatory modulation over the rising and falling portions of each sinusoidal half-cycle as the individual current components evolve. Shaded regions (I and II) indicate segments of the second LFAC cycle shown as zoomed views to highlight fiber-specific differences during depolarization (I) and hyperpolarization (II).

**Fig 8.**
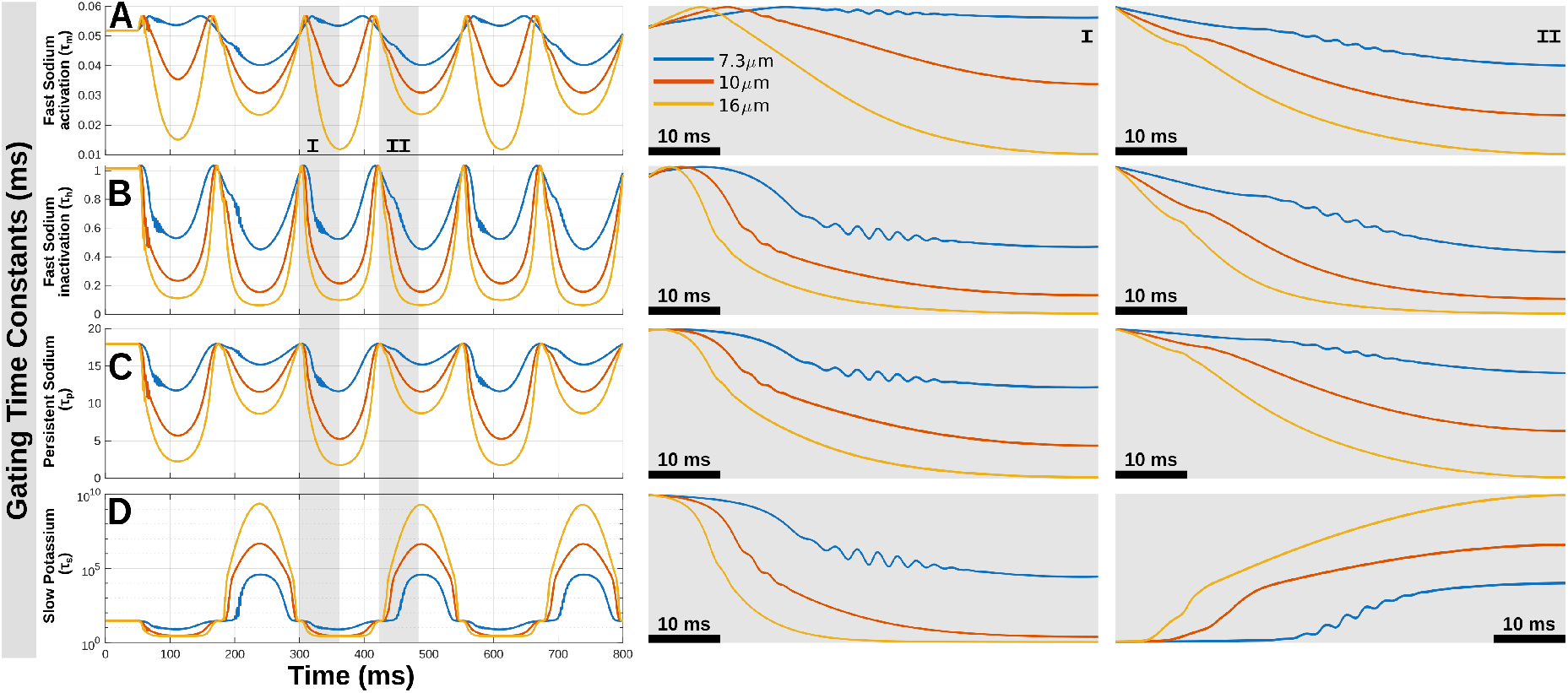
Gating time constants of nodal ion-channel states during subthreshold LFAC stimulation in the *fiber-specific* condition. Using the same stimulation protocol as in Fig 6 and 7, this figure shows the temporal evolution of the gating time constants at Node 40 for the three fibers: 7.3*µ*m (blue), 10*µ*m (orange), and 16*µ*m (yellow). (A–D) Time constants for fast Na^+^ activation (*τ_m_*), fast Na^+^ inactivation (*τ_h_*), persistent Na^+^ activation (*τ_p_*), and slow K^+^ activation (*τ_s_*), showing diameter-dependent kinetics and phase-locked modulation over the LFAC cycles. The Na-related time constants (*τ_m_*, *τ_h_*, and *τ_p_*) vary with the sinusoidal polarity and differ across diameters, with larger fibers showing smaller time constants (faster) as LFAC progress. In contrast, *τ_s_* shows the strongest phase dependence, increasing rapidly during hyperpolarization and reaching values on the order of 10^9^–10^10^ms in the 16*µ*m fiber, indicating slower slow-K^+^ kinetics relative to the depolarizing portions of the cycle. Shaded regions (I and II) indicate segments of the second LFAC cycle shown as zoomed views to highlight fiber-specific differences during depolarization (I) and hyperpolarization (II) and the influence of subthreshold oscillations on the channels’ time constants.

Correspondingly, the gating variables of fast sodium (Fig 6B for *m*, Fig 6C for *h*), persistent sodium (Fig 6D for *mp*), and slow potassium (Fig 6E for *s*) showed fiber-dependent behaviors. Subthreshold oscillations were reflected in the fast Na^+^ gating variables (*m* and *h*), indicating that the oscillatory behavior is coupled to the fast Na^+^ activation/inactivation subsystem. Across diameters, large fibers exhibit more rapid state transitions, where *h* decreases more quickly during depolarization and recovers more rapidly in the subsequent phases, while the slow K^+^ activation variable *s* rises earlier and reaches a higher plateau in the 10 and 16 *µ*m fibers compared to the 7.3*µ*m fiber. This combination indicates a stronger shift toward reduced inward transient Na^+^ activation together with enhanced outward (slow K^+^) stabilization in larger fibers over the LFAC cycle.

To assess the effective inward current flow from sodium channels, we computed the fast Na^+^ activation factor *m*^3^*h* (Fig 6F) and the persistent Na^+^ activation factor *p*^3^ (Fig 6G). During the depolarizing portion of the LFAC cycle, the large fibers exhibit a decline in *m*^3^*h* as depolarization progresses, whereas the small fiber showed a comparatively stable *m*^3^*h* magnitude. The *m*^3^*h* factor rises with depolarization in all fibers but shows stronger modulation, including subthreshold oscillations prior to decline around the peak of the depolarizing phase in the larger fibers. In parallel, the persistent Na^+^ factor *p*^3^ shows no oscillations, but their magnitude values were much higher than those of fast Na^+^ factor in a fiber-dependent manner.

The other frequency results are provided in Supplementary files (Fig 1 in S2, S3, S4, S5, and S6 for 1, 2, 3, 8, and 20Hz, respectively). Across these tested frequencies, the membrane potential at Node 40 remained phase-locked to the sinusoidal LFAC waveform, with subthreshold oscillations also observed near the depolarizing portion of the LFAC cycle and most reflected in the 7.3*µ*m fiber. However, because the LFAC period changes with frequency, the oscillations occurred at different time points within the cycle (i.e., it shifted along the depolarizing half-cycle in absolute time) as frequency varied. The gating variables (*m, h, mp, s*) and gating factors (*m*^3^*h* and *mp*^3^) showed frequency- and diameter-dependent changes. Similar to 4Hz, the fast Na^+^ gating variables (*m* and *h*) reflected the subthreshold oscillations, whereas the slower variables (*mp* and *s*) evolved gradually. At lower frequencies (1, 2, and 3Hz), the 7.3*µ*m fiber showed a clear reduction in the fast Na^+^ activation factor *m*^3^*h* during the depolarizing portion of the LFAC cycle, and the slow K^+^ activation variable *s* reached a plateau level that was not observed in the 4Hz case. As frequency increased (8 and 20Hz), the *m*^3^*h* responses became more transient within each cycle, and peak *m*^3^*h* magnitudes increased with fiber diameter, with the largest values observed in the 16*µ*m fiber. Across all frequencies, the persistent Na^+^ activation factor *mp*^3^ remained diameter dependent (larger fibers showing larger magnitudes). At 20Hz, *s* remained elevated during successive cycles and did not fully return to its steady-state level within a single LFAC period.

Fig 7 decomposes the membrane current densities into individual ionic, capacitive, and leak components. During the 4 Hz LFAC depolarizing phase, the inward fast Na^+^ current (Fig 7A) increased for a short period of time and then decreases more rapidly in the larger fibers, consistent with the concurrent reduction in *m*^3^*h* and faster changes in *h*. The persistent Na^+^ current (Fig 7B) also shows diameter-dependent modulation, including a more decline around the depolarizing peak in larger fibers compared to the small fiber. During the LFAC cycle, the slow outward K^+^ current (Fig 7C) increases earlier and to larger magnitude in the 10 and 16 *µ*m fibers, mirroring the behavior of *s* for the outward stabilizing shift with increasing fiber diameter. Notably, the capacitive current (Fig 7D) captures high-frequency subthreshold oscillations, consistent with rapid voltage fluctuations in the near-threshold regime. The mean capacitive current densities were -4.18nA/cm^2^, -6.37nA/cm^2^, and -5.89nA/cm^2^ for the three fibers (7.3*µ*m, 10*µ*m, and 16*µ*m) respectively. In contrast to the voltage-gated components, the linear leak current (Fig 7E) followed the LFAC-driven *V_m_*(shown in Fig 6A) response and there wasn’t any phase-dependent modulation, beside the subthreshold oscillations, consistent with an Ohmic passive pathway. The total membrane current (Fig 7F), which approximately sums all non-electrode transmembrane currents, was oscillating during the rising/failing portions of the sinusoidal phases. The mean values were -32.65*µ*A/cm^2^, 105.65*µ*A/cm^2^, and 375.11*µ*A/cm^2^ for the three fibers (7.3*µ*m, 10*µ*m, and 16*µ*m) respectively reflecting mostly outward current for large fibers. Furthermore, the current-density results for the other tested LFAC frequencies are provided in Supplementary files (Fig 2 in S2, S3, S4, S5, and S6 for 1, 2, 3, 8, and 20Hz, respectively). Like 4Hz case, fiber-dependent behaviors were observed in the ionic-current decomposition (fast Na^+^, persistent Na^+^, slow K^+^, capacitive, leak, and total currents). However, as frequency increase, the inward fast Na^+^ current is increased in magnitude with fiber diameter and dominated the outward component during depolarization.

Fig 8 summarizes the voltage-dependent gating time constants for fast Na^+^ activation (Fig 8A *τ_m_*), fast Na^+^ inactivation (Fig 8B *τ_h_*), persistent Na^+^ activation (Fig 8C *τ_p_*), and slow K^+^ activation (Fig 8D *τ_s_*) during the same conditions presented in Fig 6 and Fig 7. Because a gating time constant quantifies how rapidly a state variable transitions toward its steady-state value (smaller *τ* indicates faster kinetics), all four time constants were found to be phase locked to each half-cycle of the LFAC waveform, changing from their resting values after LFAC onset and evolving systematically across each half-cycle before returning toward baseline near the zero-crossings. The Na-related time constants (*τ_m_*, *τ_h_*, and *τ_p_*) showed consistent diameter dependence (seen in the highlighted windows), with larger fibers exhibiting smaller time constants (faster gate transitioning) than the 7.3*µ*m fiber. In contrast, *τ_s_*was found to be the strongest phase dependence, with different kinetics between the hyperpolarizing and depolarizing portions of the LFAC cycle; during the hyperpolarizing regions *τ_s_* increased by several orders of magnitude, reaching *∼* 10^9^–10^10^ms in the 16*µ*m fiber, which corresponds to an *∼* 8–9 order-of-magnitude (i.e., *∼* 10^8^–10^9^*×*) increase relative to its depolarizing-phase values (*∼* 10^0^–10^1^ms) and similarly for the other fibers with slower magnitudes. Consequently, *τ_s_* does not change symmetrically with the LFAC cycles as the other Na-related time constants. Further, the subthreshold oscillations were observed in all fibers as well in all gating time constants; with 7.3*µ*m fiber shown stronger oscillations. The gating time-constant results for the other tested LFAC frequencies are provided in Supplementary files (Fig 3 in S2, S3, S4, S5, and S6 for 1, 2, 3, 8, and 20Hz, respectively). With those frequencies, the time constants at Node 40 showed behavior consistent with the 4Hz case, with clear phase-locked modulation over the LFAC cycle and diameter-dependent differences. In general, larger fibers maintained smaller Na-related time constants (faster kinetics) compared to the 7.3*µ*m fiber, and they varied systematically with LFAC polarity except *τ_s_*, which also showed asymmetrical behavior. With increasing frequency, the separation in fast Na^+^ inactivation kinetics (*τ_h_*) between the larger fibers and the 7.3*µ*m fiber was reduced, with *τ_h_* in the larger fibers becoming closer in magnitude (became slower) to that of the small fiber.

In the Supplementary file (S7), we report the corresponding membrane potential, gating/gating-factor, current densities, and time-constant results under the *common-amplitude* protocol in which all fibers received the same 4Hz LFAC peak current set to the 7.3*µ*m subthreshold level of 230.9*µ*Ap. As expected, this protocol revealed differences in proximity to excitation across diameters and therefore alters the magnitude of the voltage response. Therefore, the observations under *fiber-specific* protocol became different for larger fibers (10*µ*m and 16*µ*m) and remain the same for 7.3*µ*m fiber. The large fibers responses to this subthreshold level of current weren’t observed to induce any subthreshold activity; there was no observed oscillatory behavior nor transient Na^+^ factors (captured by *m*^3^*h* or Na^+^ currents) to show attenuation during the depolarizing portion of the LFAC cycle. However, the gating time constants (Fig 3 in S7) show further asymmetrical behavior for all kinetics including Na-related time constants unlike the results of *fiber-specific* protocol. Taken together, the *fiber-specific* protocol’s results establish diameter-dependent channel-state evolution under matched near-threshold LFAC conditions, while the appendix results (*common-amplitude* protocol) provide complementary evidence under a fixed applied-current.

## Discussion

The findings of this study provide a possible mechanistic explanation for why extracellular sinusoidal LFAC stimulation preserves the natural recruitment order of nerve fibers during electrical stimulation using the MRG models of myelinated motor nerve fibers.

The first part of this study compared the intracellular and extracellular activation thresholds for both sinusoidal LFAC and pulse stimulation. The results showed that when current was injected intracellularly, both waveforms induced orderly fiber recruitment. In this case, the nerve response was independent of the extracellular space and primarily governed by the morphological properties of the membrane. These properties include capacitance and resistance, both of which scale with fiber diameter and influence current flow [11, 12]. As the internodal compartments of the MRG model incorporate myelin sheath capacitance, they are fiber diameter-dependent as well and would have influence current flow. The results demonstrated that the amount of current required to reach excitation was proportional to fiber diameter, as shown in Fig 2A and Fig 2B for pulse and LFAC via intracellular stimulation. This suggests that the inversion of recruitment order of myelinated motor nerve fibers is an inherent feature of extracellular stimulation rather than a consequence of waveform properties.

While several studies have shown that modifying waveform shape and parameters in extracellular stimulation can achieve orderly recruitment [3, 16, 33–37], their primary mechanism involves selectively blocking the propagation of action potentials. This allows only specific fibers or subsets of nerve fibers to be activated in an orderly manner. Unlike these methods, sinusoidal LFAC preserves orderly recruitment without requiring conduction block, suggesting an underlying mechanism that is intrinsic to the interaction between sinusoidal stimulation and nerve fiber biophysics.

With extracellular stimulation using bipolar cuff electrode, two modes of activation were produced: direct cathodic activation under the cathodic electrode and virtual activation outside the cuff (Fig 5). Direct cathodic activation started during the depolarizing phase of the sine wave, following the zero-crossing and prior to the peak time. On the other hand, virtual activation was induced and span around the maximum sinusoidal peak time. These findings align with our previous results [1], supporting the expectation of biphasic currents to induce both depolarization and hyperpolarization effects [38]. The activation thresholds for these two modes revealed some difference in recruitment order. Direct cathodic activation followed orderly recruitment with maintained strength-frequency relationship, whereas virtual activation thresholds were higher than cathodic with maintained strength-frequency relationship except at 20 Hz; where 5.7 and 7.3 *µ*m fibers required higher currents (Fig 2D). Besides those two fiber sizes, the the recruitment order was normal, small to large. The disturbance of the relationship at 20Hz during virtual activation for 5.7 and 7.3 *µ*m fibers can be attributed to stimulating at a relatively higher frequency, 20 Hz compared to 1-8Hz, which is closer to 35Hz where it was shown in our previous study [1] to be around the range predicted to not maintained the orderly recruitment order; and as frequency increases, the difference between fibers’ thresholds converges to minimal values during virtual activation.

The Supplementary Fig S8 shows the calculated activation function at the peak time of the direct cathodic activation threshold for three fibers, along with the electrode contact locations. The node-to-node distance and the number of nodes inside the cuff may contribute to orderly recruitment, but they are not the primary factors, as pulse stimulation still induces an inverse recruitment order despite similar geometric conditions. This indicates that the activation function alone does not explain why LFAC produces orderly activation. However, the activation function shows that larger fibers tend to have a wider virtual activation region, enhancing their virtual activation. These broader virtual sites span multiple nodes of Ranvier, more likely leading to the strong burst activations outside the cuff during the sinusoidal peak time and could lead to the disturbance of the strength-frequency relationship at 20Hz during virtual activation for 5.7 and 7.3 *µ*m fibers.

The second part of this study explored the hypothesis that large nerve fibers exhibit nerve accommodation more than small fibers. This was first tested by comparing electrotonus and threshold electrotonus; which under normal conditions should closely match, “parallels,” each other [18]. Our results showed that 100ms pulse stimulation (Fig 3A and Fig 3C) replicated the classical electrotonus and threshold electrotonus behaviors [17, 18, 39, 40] in the general response shapes. These results indicate that there is fiber dependency shown in the induced responses with both LFAC and pulse stimulation. With pulse stimulation and 100ms conditioning, activation thresholds were reduced during the course of conditioning with small nerve fibers being reduced more than large fibers (Fig 3C and Fig 3E). These results are in agreement with the results reported in the original formulation of the MRG models; where 10*µ*m fiber showed threshold reduction more than 14*µ*m fiber (figure 7A in [24]) using point source simulations. The accommodative effect on thresholds can be seen by the rapid decrease in excitability followed by fiber-dependent excitability decline during the course of 100ms conditioning (Fig 3C); known as slow accommodative response phase [39].

Unlike long pulse conditioning with threshold tracking, LFAC conditioning revealed a clear fiber-dependent changes in both electrotonus and threshold electrotonus (Fig 3B and Fig 3D). LFAC conditioning resulted in oscillatory depolarization and hyperpolarization phases during the course of conditioning. The electrotonus shape revealed a fiber-dependent distorted sinusoidal behaviors during the phasic transition from onset to the rising phase and around the zero-crossing, but not exactly at the zero crossing (Fig 3B). Threshold electrotonus with LFAC revealed that changes in thresholds and thus axonal excitability were both time and fiber-dependent (Fig 3D). The behavior of threshold changes started with reduction in all fibers (from the onset of LFAC at 50ms until around 75ms), but with reduced large fibers’ thresholds more than small fibers. About 50ms after onset, the behavior of thresholds’ changed and became more time and fiber dependent with large fibers requiring more than 200% of the initial thresholds to reach excitation at the maximum sinusoidal peak time. This behavior of thresholds changes was reversed back to the initial 50ms’ behavior as the depolarizing phase transitioned to hyperpolarization and specifically before the zero-crossing. Panels of Fig 4 expanded those in Fig 3F to illustrate how fibers recruitment order was modulated for each time delay of the test pulse during the single LFAC cycle. It is critically important to notice that action potentials’ initiation due to LFAC activation starts during the time range after the zero-crossing; however, in these threshold tracking tests, the induced action potential is synchronized to the testing pulse. A Supplementary video (S1 Video) illustrates the temporal evolution of these threshold curves across the LFAC cycle. These temporal observations along with the spatial-temporal behaviors (Fig 5) suggest that LFAC activation occurs at an equivalent time range of when higher current is needed to activate large fibers with the tracking pulse stimulation.

This accommodative effect was more defined with LFAC conditioning; where large fibers show reduced excitability (requiring more current to activate) more than small fibers, and more likely to contribute to the cathodic inverse strength-frequency relationship to achieve orderly fiber recruitment with LFAC stimulation. The cathodic strength-frequency relationship in this study and with our prior study [1] agree with the classical findings reported by Hill’s and others [41, 42]. Those classical studies stimulating frog motor nerves with sinusoidal AC frequencies below 50 Hz revealed an inverse relationship between threshold and frequency [41, 42]. Based on Hill’s accommodation theory [41, 43], those studies explored the accommodation mechanisms and axonal proprieties influencing the strength-frequency relationship, and our results agree with the inverse strength-frequency relationship. Hill et al. also noted that, at frequencies below the optimum, an AC waveform initiated near phase 0 behaves as a slowly increasing stimulus that allows the threshold to rise (i.e., promotes accommodation), whereas initiation near the crest (peak) behaves more like a constant-current step [41]. Our LFAC simulations provide a channel-level correlate of this effect (see below), showing that LFAC slow depolarization drives progressive accommodation effect in all fibers. Consistent with Hill’s description, subthreshold oscillations and action potential initiation in our simulations occurred after LFAC onset during the rising portion of the depolarizing phase, and the timing of this initiation shifted in absolute time as the LFAC period changed across our frequency range. Furthermore, we observed that initiating the sinusoid at a zero-crossing was critical for maintaining subthreshold dynamics, since any non-zero onset could introduce an immediate action potential firing, consistent with Hill’s observation that initiation near the crest (peak) approximates a step-like (DC-like) stimulus and alters the excitation condition [41].

Thus, our results of LFAC threshold electrotonus, Fig 3D, suggest that membrane accommodation could lead to orderly of fibers recruitment. This has been suggested by other studies utilizing threshold electrotonus; where the median nerve afferents, large fibers, accommodate to the long subthreshold conditioning pulse more than the sural nerve afferents, smaller fibers [20, 21]. Studies also showed that large cutaneous fibers accommodate significantly more to slowly increasing current ramp than small cutaneous fibers [22]. Our results are in agreement with those found in motoneurons, where fast twitch motor units, innervated by large motoneurons, displayed accommodation to current ramps significantly more than slow twitch motor units, innervated by small motoneurons [19]. Furthermore, in the human median nerve, it was demonstrated that large fibers appeared to accommodate to ramp prepulse more than small fibers utilizing conduction velocity measurement of Electromyography [23]. The common mechanisms underlying those responses have been attributed primarily to the dynamics of voltage-gated ion channels. Specifically, sodium channels transition to their inactivated state, while potassium channels activate, reducing the probability of action potential firing during slow changes in membrane potential [18]. In Hill’s formulation, AC excitation reflects an interaction between a time-varying local excitatory disturbance and a time-varying threshold that captures accommodation [41]. With this view, our subthreshold LFAC dynamic analysis shows that larger fibers enter an accommodated state during the depolarizing half-cycle, characterized by reduced fast-Na^+^ availability (lower *m*^3^*h*) and enhanced slow-K^+^ activation (Fig 6E and F), thereby increasing excitation thresholds. Thus, for LFAC with small-angle approximation, a linear increase can be assumed during phase transitions near zero crossings, resembling a slowly rising waveform. This behavior would imply that the LFAC waveform can be analogous to a ramp-like or slowly increasing waveform, inducing subthreshold accommodation as shown in Fig 3D. Therefore, the sinusoidal LFAC’s slow alternation would then allow the membrane to accommodate to sustained depolarization and hyperpolarization, requiring higher current thresholds to overcome this accommodation effect at low stimulation frequencies in a fiber-dependent manner leading to orderly recruitment.

In our second analysis for accommodation mechanism, we observed membrane potential oscillations during the depolarizing portion of the LFAC cycle prior to action potential initiation. In Fig 5A, subthreshold oscillations appear before spiking, and at threshold the action potential is initiated during these oscillatory fluctuations (Fig 5C, Node 40 lower trace; Fig 5F, direct cathodic activation at Node 40). This behavior suggests that membrane potential oscillations enhance membrane ability to fire APs at a depolarizing level that is otherwise subthreshold. From Fig 6A, the temporal structure of the oscillations relative to the LFAC waveform suggests that LFAC acts as a slow conditioning drive that brings the membrane into a regime where intrinsic oscillatory dynamics can arise and influence spike triggering. Dynamically, our results under the *fiber-specific* subthreshold protocol reflect subthreshold oscillations in the fast Na^+^ gating variables *m* and *h* (Fig 6B and Fig 6C) and in the capacitive current (Fig 7D); suggesting that the oscillations are coupled to the rapid Na^+^ gating and rapid voltage fluctuations. In contrast, the persistent Na^+^ and slow K^+^ activation variables (*mp* and *s*) change slowly (Fig 6D and Fig 6E) and lack subthreshold oscillations.

The subthreshold oscillations observed were not simply a direct reflection of the LFAC stimulation frequency. Instead, they appear as higher frequency fluctuations superimposed on the LFAC-driven polarization, more likely the intrinsic near-threshold responses arising from the interaction of passive membrane filtering and fast active conductances [44]. The early observations of Erlanger et al. during myelinated axons (dog phrenic and frog phalangeal nerves) excitation reflected on the presence of subthreshold oscillations and indicated that they were more spontaneous to raise fiber’s excitation state to threshold [45]. Tasaki et al. showed that low frequency alternating currents (50-3000Hz) gradually modify the state of the nerve membrane before threshold excitation is reached [46]. Their experiments on the toad motor nerve revealed that AC stimulation accumulates excitability over time with membrane potential fluctuations, shifting the excitation threshold in a frequency-dependent manner [46]. Thus, our results suggest subthreshold oscillations as a precursor to AP initiation that could increase the probability of AP generation when they transiently elevate excitability level.

We found that ion channels dynamics were all fiber- and frequency-dependent; where larger fibers were accommodating to LFAC more than small fibers. During 4Hz LFAC stimulation, the main accommodation characteristic is the reduction in the fast Na^+^ activation factor *m*^3^*h* of the large fibers during the depolarizing portion of the LFAC cycle (Fig 6F), accompanied by attenuation of the inward fast Na^+^ current (Fig 7A). This decline in *m*^3^*h* suggests that the transient Na^+^ channel population becomes less available for activation (i.e., a shift toward a more inactivated state), while in parallel the slow K^+^ activation increases more rapidly and reaches a higher plateau in larger fibers (Fig 6E), consistent with an outward stabilizing influence. In contrast, the small fiber maintains a sustained transient Na^+^ activation and shows oscillatory fluctuations in *m* and *h* (Fig 6B and Fig 6C), producing an oscillatory modulation of inward Na^+^ current (Fig 7A) that could facilitate spike triggering at lower current levels. When decreasing LFAC frequency to 1Hz (Supplementary file S2), the small fiber started to reflect accommodation via similar mechanisms (like large fibers during 4Hz LFAC stimulation: attenuated *m*^3^*h* and fast Na^+^ current with an increased K^+^ activation) and accommodation of large fibers became stronger (much lower values for *m*^3^*h* and inward fast Na^+^ current than during 4Hz LFAC stimulation). As frequency increased up to 20Hz (Supplementary file S6), large fibers started to be less affected by accommodation, and their state dynamics became similar to those of small fibers during 4Hz LFAC stimulation. This combination of reduced effective transient Na^+^ drive with enhanced slow K^+^ activation agrees with the classical definition of the accommodation state [18, 41] that prevents action potential firing during slow depolarization at lower AC frequencies. This aligns with threshold-tracking interpretations in which time-dependent, local excitability changes reflect both passive polarization and nonlinear active components near the stimulating site [47].

Under the *common-amplitude* protocol (Supplementary file S7), all fibers receive the same LFAC peak current (set to the 7.3*µ*m subthreshold level), which indicates differences in proximity to excitation across fiber diameters. In this condition, subthreshold oscillations are induced mainly in the small fiber, whereas the larger fibers remain farther from excitation and therefore do not exhibit oscillatory behavior or near-threshold Na^+^ modulation. This is consistent with the expectation that, even under the same applied stimulus, fiber geometry and electrical properties lead to diameter-dependent polarization and effective recruitment conditions; thus, the induced subthreshold channel-state changes differ across fibers when the same stimulus amplitude is applied. Therefore, *fiber-specific* protocol establishes the mechanistic ordering under matched near-threshold conditions, while *common-amplitude* protocol illustrates how a fixed stimulus emphasizes recruitment separation and suppresses near-threshold oscillatory features in fibers that remain farther from threshold.

Our findings also suggest that the mechanisms shaping subthreshold accommodation under LFAC depend on both frequency and fiber diameter. At lower frequencies, the longer depolarizing phase enhances the contribution of nonlinear active membrane processes, inducing more fiber-dependent accommodation characteristics. As frequency increases, the within-cycle accommodation effect is reduced and the response becomes more influenced by the intrinsic temporal kinetics of the channel states, which are classically known to be diameter dependent (larger fibers have faster gating transitions and smaller fibers slower transitions). Therefore, at 20Hz LFAC stimulation the activation thresholds across diameters become closer (Supplementary file S6), due to the reduced time per half-cycle for accommodation to fully develop while kinetic differences still shape the cycle-to-cycle channel-state responses.

Furthermore, the gating time constants analysis reveals that LFAC drives fiber-dependent and phase-dependent kinetic regimes (Fig 8). Larger fibers exhibit smaller Na-related time constants over the LFAC cycle (faster), particularly for fast Na^+^ inactivation *τ_h_*, whereas the 7.3*µ*m fiber maintains comparatively longer time constants. The strong phase dependence of the slow K^+^ time constant *τ_s_*, including its large magnitude increase during hyperpolarization, suggests that slow K^+^ gating can act as a stabilizing factor over the LFAC cycle; where fast Na^+^ inactivation maximum is reached while potassium channels still active. Thus, the gating variables, current, and time constants results suggest that LFAC does not only depolarize and hyperpolarize the membrane symmetrically; rather, it drives a cycle-by-cycle channel state response that facilitates accommodation in larger fibers while allowing the oscillation-facilitated action potential initiation in smaller fibers.

## Conclusion

This study utilized computational modeling to investigate the biophysical mechanisms of orderly recruitment during extracellular sinusoidal LFAC stimulation. Using a bipolar cuff electrode in a volume conductor framework coupled to MRG myelinated axon models, we showed that the inverse recruitment order observed with extracellular stimulation is a geometric/electrical consequence of extracellular excitation and is theoretically independent of the stimulation waveform. Simulations with bipolar cuff electrodes revealed that LFAC induces two modes of activation: direct cathodic activation occurring inside the cuff and virtual activation occurring outside the cuff.

To explain why LFAC can produce size-wise recruitment, we focused on subthreshold accommodation and near-threshold dynamics. Threshold tracking and dynamic-state analyses demonstrated that larger fibers accommodate more under slow depolarizing current, resulting in increased activation thresholds relative to smaller fibers. Under LFAC, this accommodation effect resulted in reduction in the effective transient sodium drive (reduced fast Na^+^ activation factor) along with a buildup of outward stabilization by slow K^+^ activation, producing a state in which fast Na^+^ channels become closed while slow K^+^ channels remain activated/open. In addition, LFAC induced subthreshold oscillations that promoted action potential initiation during gradual depolarization via coupled passive membrane filtering and fast voltage-gated conductances. From the extracellular stimulation prospective, this provides a possible mechanistic basis for orderly recruitment under LFAC, in which slowly depolarization allows accommodation to develop and preferentially raises the activation thresholds of large fibers compared to small fibers.

Our findings suggest subthreshold accommodation as a contributor to LFAC orderly recruitment and suggest that LFAC can leverage intrinsic membrane dynamics to achieve controlled recruitment without requiring complex selective-block paradigms. This mechanistic understanding strengthens the rationale for LFAC as a neuromodulation strategy with potential utility in clinical peripheral nerve stimulation and neuroprosthetic applications where selective and physiologically ordered activation is desirable.

## Limitations

In this study, a bipolar cuff electrode was used to investigate extracellular stimulation mechanisms under LFAC. Our preliminary work on electrode influence on LFAC efficiency suggest that stimulation configuration (or electrode configuration) have a role in determining stimulation efficiency and the ability to maintain orderly recruitment. Therefore, a comprehensive mechanistic analysis should consider electrode configurations. It is important to note that the findings presented in this study are specific to the bipolar configuration and the parameter settings described in the methods section.

Another limitation of this study is that all mechanistic analyses were conducted using a selected range of frequencies (1-20Hz). Although this frequency range was selected based on our prior findings that fiber-dependent recruitment properties would be preserved below 35Hz, extending the analysis to additional frequencies could provide further insights into the interaction between stimulation frequency and ionic channel dynamics. In particular, examining how frequency modulates large fiber accommodation and subthreshold oscillations could improve our understanding of the conditions under which LFAC maintains orderly recruitment.

Furthermore, in this study, one electrode contact was consistently positioned at the center of the fiber models, aligned with the center of a node of Ranvier. While this approach provided controlled implementation, variations in internodal distance could introduce a range of possible activation outcomes. The relative positioning of the electrode contacts with respect to the nodes of Ranvier is known to influence stimulation efficiency. Therefore, future studies could expand the variability of contact placement based on fiber-specific internodal distances to provide a more detailed and comprehensive prediction of activation behaviors.

Lastly, virtual activation thresholds were quantified and discussed using the activation function and the predicted frequency-dependent trend observed up to 20Hz. However, the biophysical origin of virtual activation and its modulation with frequency were not investigated in detail in this study. Further work is needed to determine how LFAC frequency and propagation dynamics outside the cuff interact to shape virtual activation thresholds and recruitment behavior.

## Supporting information

**S1 Video. Threshold Tracking under subthreshold 4Hz LFAC stimulation.** Threshold tracking results for subthreshold accommodation testing using a bipolar cuff electrode. The right panel shows the LFAC conditioning waveform (one 4 Hz cycle) and the delayed 1ms testing pulse used to test excitability. For each fiber diameter, the LFAC conditioning amplitude was set to 40% of that fiber’s LFAC activation threshold. The testing pulse polarity was matched to the instantaneous LFAC polarity at each delay time point. The experimental setup is illustrated in main Fig 1C. The left panel shows the corresponding threshold-fiber diameter relationship at each delay time point, showing how the required 1 ms pulse amplitude changes across the LFAC cycle under subthreshold conditioning. A static summary of these threshold-diameter curves is provided in main Fig 4.

**S2. Subthreshold 1Hz LFAC** stimulation during the *fiber-specific* condition results.

**S3. Subthreshold 2Hz LFAC** stimulation during the *fiber-specific* condition results

**S4. Subthreshold 3Hz LFAC** stimulation during the *fiber-specific* condition results.

**S5. Subthreshold 8Hz LFAC** stimulation during the *fiber-specific* condition results.

**S6. Subthreshold 20Hz LFAC** stimulation during the *fiber-specific* condition results.

**S7. Subthreshold 4Hz LFAC stimulation during the *common-amplitude* condition results.**

**S8.**
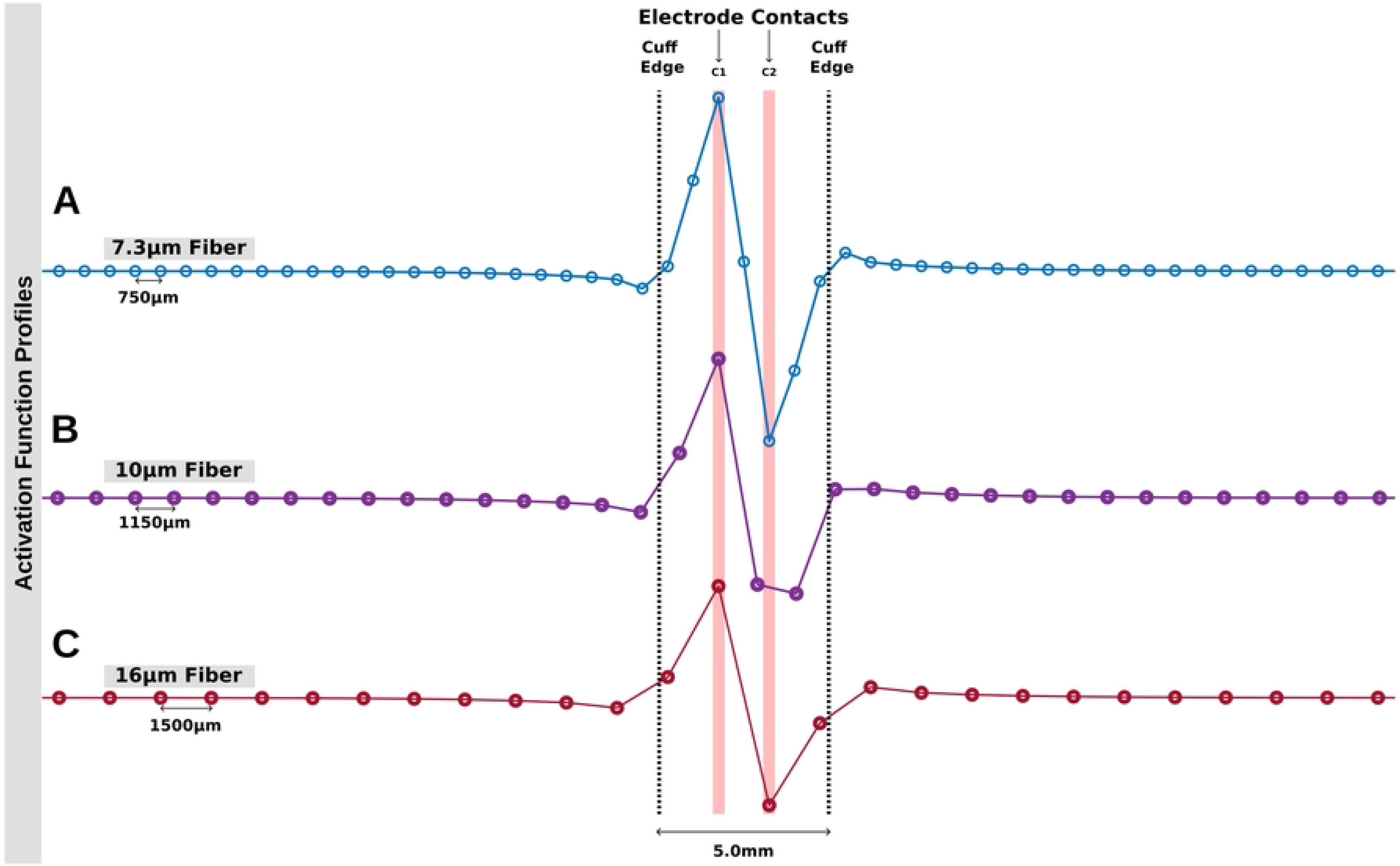
Activation function profiles for three fiber diameters. 7.3*µ*m (A), 10*µ*m (B), and 16*µ*m (C) within the same bipoar cuff electrode configuration, each at their respective LFAC direct cathodic activation thresholds at the cathodic peak time. The larger fibers have larger internodal distances, resulting in fewer nodes being positioned within the electrode region to facilitate direct cathodic activation. In contrast, virtual activation regions are broader. The electrode contacts (C1 and C2) are highlighted in red, and the cuff electrode edges are indicated by dashed lines.

## Acknowledgments

The authors would like to thank Dr. M. Ryne Horn and Mr. Nathaniel Lazorchak for providing their initial scripts/models and discussing the experimental results.

